# Assessment of *Blighia sapida* on Cholinergic and Antioxidant Enzymes; Possible Use of the Plant Stem-Bark Extract as a Biological Pest Controlling Agent

**DOI:** 10.1101/2023.05.30.542835

**Authors:** M.B. Adekola, O.V. Oriyomi

## Abstract

The harmful effects of synthetic pesticides include neurological, behavioural dysfunctions, hormonal imbalances, and water pollution. Hydro-alcohol extract of the stem bark of *B. sapida* was studied for pesticidal effects on Glutathione S-transferases (GST) and acetylcholinesterase (AChE) using a rat model. Various concentrations of the extract were administered to six different groups, of three male and three female groups of rats (50 mg/kg, 100 mg/kg and 150 mg/kg bwt. respectively), two synthetic 10% (w/v) groups and a control group. Blood plasma, liver, and brain were obtained at the end of 28 days sub-acute test, from the Wistar rats for biochemical assay.

The results showed that there was a significant decrease (P < 0.05) in acetylcholinesterase (AChE) activities in the brain of female rats while that of males was not significant (P > 0.05) compared to control. Also in GST, a significant increase (P < 0.05) in all the doses in liver but only at 100 mg/kg and 150 mg/kg in plasma of female rats, was observed compared to control while in male rats, a significant increase in both liver and plasma at 100 mg/kg and 150 mg/kg but not significant at 50 mg/kg was observed. The non-significant difference was observed in alkaline phosphatase (ALP) aspartate aminotransferase (AST), alanine aminotransferase (ALT) and total protein (TP) activities in both sexes at 50 mg/kg compared to control.

It was concluded that hydro-alcohol extract of *B. sapida* significantly reduced the levels of AChE and GST activities at higher and lower doses respectively. This property of the plant could be exploited in the formulation of agents useful in pest control.

## 1. Introduction

Pesticides are compounds formulated to destroy, control or prevent pests that can damage agricultural products. Since ancient times, man has engaged in agriculture to meet his food needs and has used pesticides to prevent pre- and post-harvest losses of crop due to pest infestation (Saroj et al., 2020). Pesticides commonly consumed in agricultural production and protection include insecticides (7.50%), fungicides and herbicides, (12.06%), herbicides (25.1%) and algaecides, nematicides and acaricides (53.84%) of the total global pesticides production (Koul et al., 2008; Laxmishree and Singh, 2017). Commonest are the organochlorines, organophosphates and carbamates-available as insect repellents (e.g. diethyltoluamide (DEET), carbofuran and aldicarb (Casida and Durkin, 2013; Chowański et al., 2014; Enyiukwu et al., 2016), Fumigants (e.g. 1, 2–dibromoethane, chlorophenols, methyl bromide, diquat) (Erhunmwunse *et al*., 2012) and insecticides (e.g. triazine) (Gupta and Misra, 2017). Synthetic organic pesticides are extremely toxic, due to their persistence in the environment and absorption by human or animal (e.g. cyanide aluminium-phosphate and methyl-bromide (Laxmishree and Singh, 2017).

Synthetic pesticides also demonstrate harmful effects such as neurological, behavioural dysfunctions, hormonal imbalances, kidney and liver disorders, genotoxicity and many others (Zibaee, 2011). Synthetic pesticides constitute major causes of water pollution, air and water contamination (Ravindran, 2016). Because of their non-biodegradability coupled with persistent nature, they tend to accumulate in the environment, causing ecosystem disturbances. The environmental-implication of synthetic pesticides includes but not limited to environmental-pollution, reduction in biodiversity and destruction-of-marine -life, rapid development-of-resistance-in-pests and adverse-effects-on-non-target organisms (Malhat et al., 2018). Taken into consideration, the negative effects of synthetic pesticides on the environment and non-target organisms have led to a general belief that natural compounds are better alternatives (Hassan and Gokce, 2014; Chowdhary et al., 2018).

Understanding the mode of action of pesticides is important as it reveals whether an insecticide is toxic to non-target organisms such as fish, birds and mammals or not. Important insects like bees that pollinate plants can be killed by insecticide and this would result in a reduction in crop yields (Garcia et al., 2012). Acetylcholinesterase is a key enzyme that terminates nerve impulses by catalyzing the hydrolysis of a neurotransmitter, acetylcholine into acetic acid and choline in the nervous system of various organisms. Acetylcholinesterase inhibitors bind to the enzyme, cholinesterase and prevent breaking down of the neurotransmitter thus, enhancing the accumulation of acetylcholine at the synapses causing rapid twitching of voluntary muscles and paralysis (Umar et al., 2015). Also, glutathione S-transferases play an important role in insecticide resistance getting involved in the metabolism of organophosphorus and organochlorine compounds (Zazali, 2016). GSTs are the mainly cytosolic enzymes that catalyze the conjugation of electrophile molecules with reduced glutathione (GSH), making potentially toxic substances become more water-soluble and generally less toxic (Dobritzsch *et al*., 2020). The resulting GSH conjugated compounds are eliminated from cells as reported by Lee et al. (2015).

Inhibitors of acetylcholinesterase include but not limited to organophosphates and the carbamates (Zhang et al., 2017). Some insecticides have negative effects on the nervous, renal, respiratory and reproductive systems of insects. Insecticides (organochlorines, organophosphates and carbamates) are designed to attack an insect’s nervous system and capable of producing acute and chronic neurotoxic effects in mammals (Pavela, 2016).

Botanical pesticides exhibit a number of merits over synthetic products (Dimetry, 2014). The environmental safety of botanical pesticides is one of their main positive aspects. The active substances of botanical pesticides are very friendly to many non-target organisms. Presently, biopesticides have contributed to the minimization of environmental and health problems associated with the application of some synthetic products (Pavela, 2016). Botanical pesticides or botanicals are one of the alternatives to conventional pesticides. Botanicals are secondary metabolites that are present in the plants and are considered safe. They play an important part in agricultural pest management (Isman, 2006). Therefore, there is a need to investigate more plants for their insecticidal activities since they are ecofriendly and have little or no effects on non-target organisms to minimize the effects of synthetic pesticides on both the environment and non-target species.

## 2. Methods

### 2.1. Sample collection

Fresh stem bark peelings of *Blighia sapida* were obtained at Fadama farm location (7.23443 °N and 3.43617 °E) at the Federal University of Agriculture Abeokuta, Ogun State Nigeria. The plant was identified and authenticated at the department of pure and applied botany, Federal University of Agriculture Abeokuta.

### 2.2. Preparation of Hydro-alcohol extract of plant materials

*B. sapida* stem bark was air-dried for two weeks at 25 ± 2 °C and ground into powder by electrical Grinding Machine. The powder material (1066.71g) was macerated in 70 % (v/v) ethanol/water for 72 hours at room temperature using the method of (Handa *et al*., 2008). The resulting suspension was filtered and strained with a muslin cloth. The filtrates were pooled together and concentrated with a rotary evaporator at 40 °C to yield a residue termed ethanol extract (EE). The resulting extract was weighed, labeled and stored in the desiccator until required for further analysis.

### 2.3. Experimental Animals

Wistar rats of both sexes, weighing between 150 and 220 g used were obtained from the Department of Anatomy, University of Ibadan, Ibadan, Nigeria. The animals were acclimatized to the laboratory conditions for two weeks in the Animal House and fed with standard pellet with free access to water. The principle of laboratory animal care (NIH publication No. 85–23) guidelines and procedures were followed in the study (NIH publication revised, 1985). Animal handling and care complied with international laboratory animal use and care guidelines.

#### 2.3.1. Animal grouping

The male and female Wistar rats (54) were randomly divided (Adekola et al., 2020) into nine groups, and with each group contained six rats;

Group 1 – received 1.0 ml of distilled water;

Group 2 – Male rat received Rambo pesticide 10% (w/v)

Group 3 – Female rat received Rambo pesticide 10% (w/v)

Group 4 – Male rat received 50 mg/kg EE

Group 5 – Male rat received 100 mg/kg EE

Group 6 – Male rat received 150 mg/kg EE

Group 7 – Female rat received 50 mg/kg EE

Group 8 – Female rat received 100 mg/kg EE

Group 9 – Female rat received 150 mg/kg EE

The rats were treated by gavage administration of the extract and locally produced insecticide ‘Rambo’ containing 0.6% permethrin (synthetic pesticide) every other day for 28 days. Otitoju and Onwurah (2006) reported that 10% of Rambo is relatively safe to non-target organisms. On day 28, all the animals were fasted overnight, sacrificed and blood obtained through cardiac puncture. Collected blood together with excised liver and brain were kept in labeled heparinized bottles for the estimation of biochemical parameters.

#### 2.3.2. Preparation of blood plasma

Blood plasma was prepared according to the standard procedure of Chawla (1999) as reported by Adekola *et al*. (2020). Typically, blood samples collected in heparinized bottles were centrifuged at 3000 rpm for 10 min in a Table Centrifuge (Model 90–2) at 25 °C. The plasma was collected in sterile bottles using sterile Pasteur pipettes and kept for biochemical analyses.

#### 2.3.3. Preparation of liver and brain homogenates

The liver and brain homogenates were prepared as described by Babalola and Areola (2010). One gram (1 g) of tissue was homogenized with 100 mM of phosphate buffer, pH 7.2, to produce 10% (w/v) homogenates with pestle and mortar. The homogenates were carefully transferred into centrifuge tubes and volumes adjusted to 10 ml. This was centrifuged at 4000 rpm for 30 min in a Table Centrifuge (Model 90–2) at 25 °C. The supernatants were collected into clean sample-bottles, labeled and kept in the freezer for assay of biomarker enzymes.

#### 2.3.4. Determination of biochemical parameters

The concentrations of the biochemical parameters were assayed according to the methods described by Lowry et al. (1951) for total protein, albumin by Pinnell and Northam (1978), total bilirubin, direct bilirubin by Jendrassik and Grof (1938), activities of ALT and AST as described by Reitman and Frankel (1957) while ALP was according to the method of Klein and Kaufman (1967), Habig et al (1974) for GST and Voss and Sachsse (1970) for AChE.

### 2.4. Statistical analysis

The data obtained were analyzed using one-way analysis of variance (ANOVA) followed by Tukey–kramer multiple comparisons test using the software Graph pad Prism 5. The statistical significance set at p < 0.05. Values were expressed as mean ± standard error of the mean (SEM).

## 3. Results and Discussion

The results of the study showed that *B. sapida* exhibited a pesticidal effect in Wister rats administered with graded concentrations of the extract and synthetic pesticide (Rambo). Dose-dependent changes in AChE and GST activities in the tissues of both male and female rats are presented in Tables 1.1 and 1.2 for the control and experimental groups. The alterations in activities of other biomarkers such as ALT, total protein, total and direct bilirubin were also observed for both male and female experimental rats that were administered with 50, 100 and 150 mg/kg extract and 10% (w/v) synthetic pesticide (Rambo) as shown in Tables 1.1 and 1.2. Biomarkers are commonly used to evaluate the effect of exposure to xenobiotics (Anantha et al., 2012). This is because of their ability to reflect potentially damaging effects caused by foreign compounds. Biochemical parameters reveal effects at the sub-cellular level before apparent at higher levels of biological organization (Otitoju and Onwurah, 2007). Hydro-alcohol extract of *B. sapida* elicited a pesticidal effect on metabolizing enzymes (GST and AChE). The AChE (in the brains) and GST (in liver and plasma) activities were examined in the experimental animals administered with synthetic pesticide and Hydro-alcohol extract of *B. sapida*.

**Table 1.1:** Effect of Hydro-alcohol Extract of *Blighia sapida* on Biochemical Parameters for Female Rats.

| Groups | Control | Synthetic | Extract | Extract | Extract |
| --- | --- | --- | --- | --- | --- |
| Parameters | 0 mg/kg | 10 % (w/v) | 50 mg/kg | 100 mg/kg | 150 mg/kg |
| AChE (U/L) | 224.43 ± 4.47 | 162.93 ± 16.22* | 117.23 ± 1.50*** | 88.51 ± 5.91*** | 88.59 ± 3.37*** |
| GST Liver (μmol/min) | 134.37 ± 7.42 | 608.80 ± 14.35*** | 242.48 ± 36.10*** | 278.36 ± 28.20*** | 329.05 ± 8.10*** |
| ALP (U/L) | 0.022 ± 0.002 | 0.028 ± 0.006 | 0.017 ± 0.001 | 0.024 ± 0.007 | 0.028 ± 0.006 |
| AST (U/L) | 290.87 ± 5.83 | 302.06 ± 14.07 | 297.84 ± 9.75 | 302.36 ± 3.33 | 314.17 ± 4.60 |
| ALT (U/L) | 8.79 ± 1.09 | 23.49 ± 2.56*** | 15.74 ± 1.00 | 16.38 ± 2.08* | 24.22 ± 2.98*** |
| TP Liver (g/L) | 126.26 ± 0.75 | 146.43 ± 11.46 | 133.21 ± 8.09 | 178.35 ± 38.93 | 214.28 ± 47.61*** |
| TP Plasma (g/L) | 71.27 ± 1.23 | 142.10 ± 7.84*** | 53.81 ± 1.18 | 59.36 ± 2.29 | 85.35 ± 1.69 |
| Albumin (g/L) | 45.54 ± 1.48 | 17.26 ± 0.21*** | 17.53 ± 0.96*** | 21.11 ± 0.24*** | 21.12 ± 0.45*** |
| D. Bilirubin (mg/dl) | 10.47 ± 0.53 | 1.06 ± 0.05*** | 1.61 ± 0.10*** | 2.04 ± 0.26*** | 2.08 ± 0.21*** |
| T. Bilirubin (mg/dl) | 8.99 ± 0.69 | 1.74 ± 0.05*** | 1.96 ± 0.05*** | 2.23 ± 0.09*** | 2.79 ± 0.57*** |
| GST Plasma (μmol/min) | 500.44 ± 7.21 | 777.60 ± 19.42*** | 566.67 ± 1.56 | 598.09 ± 27.68* | 605.38 ± 18.40** |

**n=5; Mean ± standard error of mean, \*=significantly different from control at p < 0.05**

**Table 1.2:**
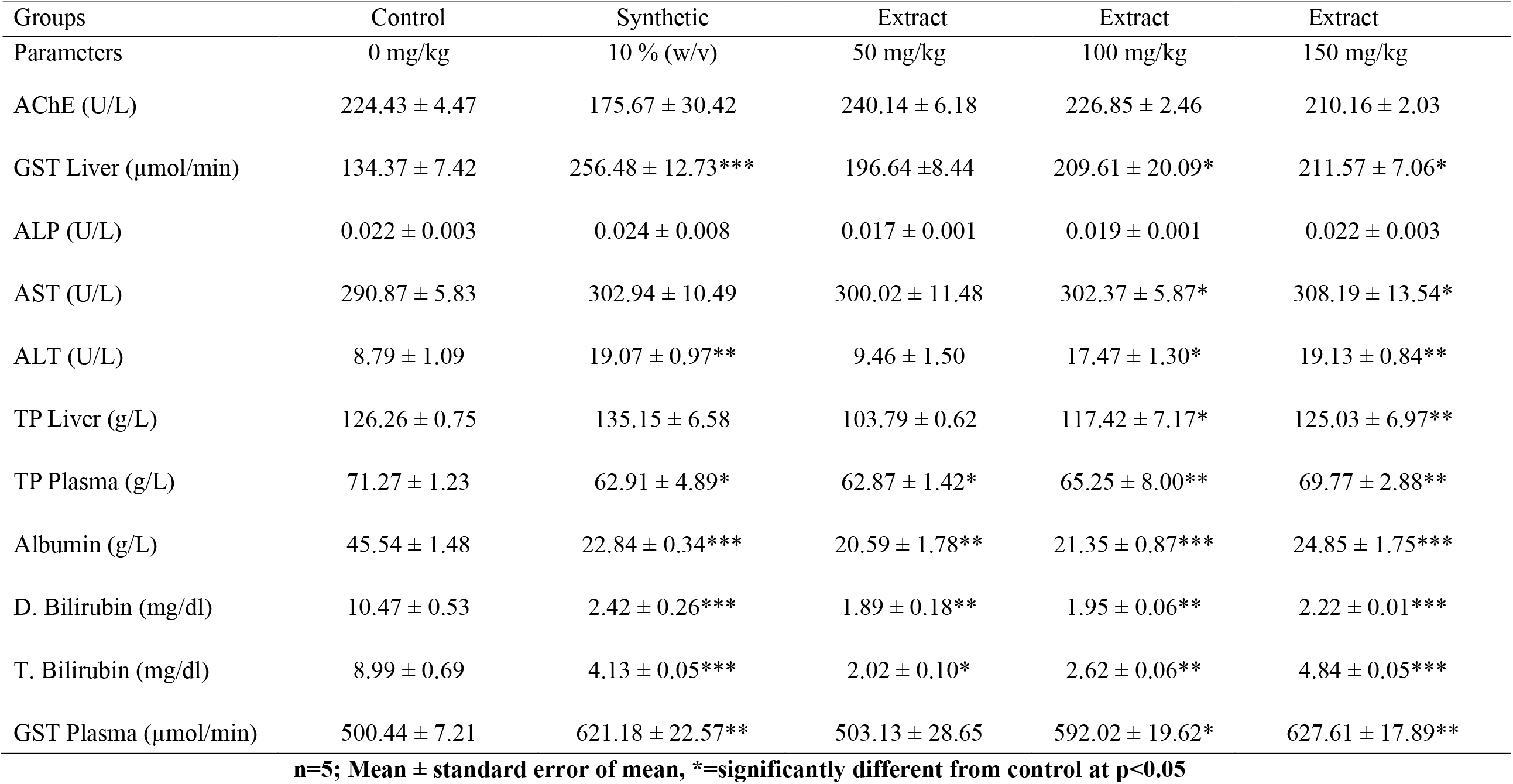
Effect of Hydro-alcohol Extract of *Blighia sapida* on Biochemical Parameters for Male Rats.

Acetylcholinesterase activity showed that, there was no significant difference (P > 0.05) for male groups compared to the control group (Table 1.2). Whereas, AChE activity was significantly different at (P < 0.05) for female groups compared to the control group (Table 1.1), the activity displayed a decreased level in a dose-dependent manner. Thus, the reduction was an indication of biopesticidal effects, leading to high levels of acetylcholine in nervous tissue hence, interruption of the proper function of the nervous system which collaborates the findings reported by (Vences-Mejia et al., 2006; Malhat et al., 2018). Glutathione S-transferases (GST) are the main cytosolic enzymes that catalyze the conjugation of electrophile molecules with reduced glutathione (GSH), which potentially play an important role in the detoxification of xenobiotics. The xenobiotics are converted to more water-soluble (hydrophilic), less toxic and easily excreted from the system.

The results of the liver and plasma GST were found to be significantly different at (P < 0.05) compared to the control group for both female and male animals, which was in agreement with Arbind et al. (2014) as reported by Polat et al. (2017). It was observed that the plant extract reduced the levels of AChE in a dose-dependent manner in male and female Wister rats. Also, at lower concentration (50 mg/kg) of the extract, the levels of GST activities in both plasma and liver homogenate, for male and female rats were lower than what was observed in 10% of Rambo (synthetic pesticide). This is an indication that the plant extract may likely reduce the resistance by GST during application.

The assessment of the activities of enzymes such as ALT, AST and ALP provides information on the liver function. ALT is a cytosolic based while AST is domicile in mitochondrial, both of which are found at a moderate concentration in the liver. Damage to the liver has been reported as the major cause of high levels of these enzymes in the plasma. Alkaline phosphatase is a membrane-bound enzyme and it plays a critical role in the transportation of metabolites across the cell membrane, conditions such as liver damage and bone disease lead to its alteration which affects the transport of metabolites and membrane permeability (Simon et al., 2019). The ethanol extract of the stem bark of *B. sapida* displayed no significant difference in the ALP for male and female rats as well as AST levels for female at (p > 0.05) while a dose-dependent increase in the activities of ALT for female and male as well as AST for male rats is an indication that the extract will be effective without any side effect in controlling pests at the lowest dose (50 mg/kg) used in the study. The results of ALP implied that phytoconstituents in the plant do not cause any damage to the membrane, hence transportation of metabolite remains intact.

The liver is mainly involved in the synthesis of plasma protein, damage to which causes a reduction in the protein concentration (Adeoti et al., 2017). A decrease in the level of albumin as well as total protein has been associated with damage to the liver function, caused by toxicity (Simon et al., 2019). In this study, the administration of ethanol extract of the stem bark of *B. sapida* caused no significant difference in the protein concentration when the experimental groups were compared with control group except in the highest dose (150 mg/kg) body weight where the total protein concentration in the liver was significantly higher than control while 10% permethrin (Rambo) group displayed a significant increase in plasma protein concentration for female rats. In the case of male rats, almost all the groups (extract-treated and Rambo groups) produced significant difference at (P < 0.05) compared to control group in the liver and plasma protein concentrations with exception of groups treated with Rambo and 50 mg/kg for liver total protein (Table 4.1 and Table 4.2). However, a significant difference in albumin level was observed in all the experimental groups (extract-treated and Rambo groups) compared to the control group. Also, there was a significant difference in the levels of both total and direct bilirubin in all the experimental groups compared to the control group.

The results of this study indicated that administration of the plant extract as biopesticide may not alter the function of the liver as displayed by liver marker enzymes, ALT, AST and ALP most especially at the doses of 50 and 100 mg/kg body weight for both male and female rats, since these enzymes are sensitive markers that indicate any damage done to the liver, being the salient organ in the detoxification and elimination of xenobiotic (pesticides). However, the significant difference observed in protein concentration may be as a result of alteration in protein synthesis in the animals.

Interestingly, there were concentrations dependent decrease in AChE and an increase in GST activities for both male and female experimental animals, which are indications of insecticidal effects of the study plant. These enzymes (AChE and GST), play a crucial role as indicators in the detection of toxicants. The work corroborates the report of Polat et al. (2017) on a decrease in the activities of AChE in animals exposed to organophosphate insecticide, a decrease in AChE activity in the frontal cortex of the rats exposed to metabolites of aspartame and the inhibition of AChE activity in rat hippocampus aspartame metabolites. The results of this study imply that bioactive compounds present in the study plant may cause a reduction in the activities of enzymes involved in the metabolism of foreign substances (pesticides) thereby leading to consequences such as tremor, reduction in the rate of survival and death. The bioactive compounds in the plant may also serve as growth retardant, reduction in the rate of reproduction, loss of weight in larva, pupa and adult.

## 4. Conclusion

The Hydro-alcohol extract of *B. sapida* significantly lowered the level of acetylcholinesterase and glutathione-s-transferases activities. This suggested *B. sapida* as having neurodegenerative ability peculiar to pesticide agents. Hence, the use of the plant extract as a repellent and for mitigation of pest control in agriculture can be adopted, which serve as an alternative, safe and ecofriendly to synthetic pesticides, the pesticidal application can be carried out both on stores and farmlands, either in powder or extract form.

## Contributors

All authors contributed to this research.

